# Reference intervals of spot urine copper excretion in preschool children and potential application in pre-symptomatic screening of Wilson’s disease

**DOI:** 10.1101/719369

**Authors:** Nelson Leung-sang Tang, Joannie Hui, Dan Huang, Man Fung Tang, Xingyan Wang, Junyi Wu, Iris HS Chan, Ting Fan Leung

## Abstract

**Background:** With spot urine collected from a large control sample of preschool children (aged 3-7 years), reference range of spot urine copper excretion indexes and their biological variation were defined.

**Methods:** In order to investigate their test performance in screening of Wilson disease in this age group, multiple spot urine samples from 6 WD patients diagnosed at presymptomatic stage were analysed. Cut-off values for spot urine copper concentration, copper to creatinine ratio and copper to osmolality ratio at 0.5 µmol/L, 0.1 µmol/mmol and 0.00085 µmol/mOsmol (32 µg/L, 56 µg/g creatinine and 0.054 µg/mOsmol, respectively, in conventional units) have potential application in differentiation of WD patients.

**Results:** The data provides a new insight that the inter-individual variation of spot urine copper indexes (CVg) were moderate with figures around 60% which was similar to other clinically useful urine tests, such as urine albumin excretion ratio. Spot urine copper excretion strongly correlated with both urine creatinine and osmolality. And more than 95% of data points in health preschool children fell within prediction regions by linear regression suggesting a good utility of normalisation by these 2 analytes. Receiver operator curve (ROC) showed that copper to osmolality ratio was the best index with an area under curve (AUC) greater than 0.98.

**Conclusions:** Based on the data, a new WD screening time window targeting preschool children is proposed. Application of a bivariate screening strategy using spot urine copper concentration and urine osmolality may be useful in a population screening program for preschool children.

## 1. INTRODUCTION

Wilson’s disease (WD) is a common inherited disease of copper metabolism caused by mutations in *ATP7B* gene (OMIM: 277900). The disease incidence in the range of one in thousands is one of the highly prevalent inherited metabolic diseases (IMD), particularly in Asian populations (1,2). In the past, WD were typically diagnosed after symptoms onset due to cumulative damage to the liver (hepatic presentation) or CNS (neurologic presentation) after a prolonged period of copper accumulation and diagnosis has been made with a clinicopathologic scoring system (3,4). Nowadays, with the advance in molecular diagnosis, a clear-cut definitive diagnosis for both affected and heterozygotes can be readily made. As early diagnosis and treatment are the key determinants of prognosis, there have been attempts of screening using various biomarkers (5). Here, we studied the reference interval of spot urine copper excretion in healthy preschool children (age 3 to 7) and examined the feasibility of its application as a screening biomarker for WD.

With the help of mutation analysis for diagnosis the natural history of WD is changed. More and more young presymptomatic WD patients have been diagnosed before adolescence and treatment can be started early with the potential of preventing significant organ damage and hopefully attaining uneventful life expectancy similar to healthy individuals (6–9). Early and safe treatment of these presymptomatic WD with oral zinc therapy is effective in reducing dietary copper absorption and thus body copper load (6–9).

The benefit of early treatment stems from the ability to make early diagnosis in WD patients particularly before 10-years-of-age. This represents a wide (a decade long) presymptomatic window period for making the early diagnosis before significant tissue damage occurs. Nature history of WD also supports this notion. First, neurological diseases have a typical onset after adolescence (2). Second, liver toxicity due to cumulative copper toxicity also commonly occurs after 10-years-of-age (8). Currently, presymptomatic patients were largely picked up by family screening after index cases had been diagnosed. While others were solely relied on clinical vigilance of the paediatricians when managing patients with unexplained symptoms and laboratory results, such as persistent elevation of liver enzymes (10). However, no biomarker suitable for large-scale case screening is available though some attempts had been reported (5).

While 24-hour urine copper is the gold standard in the assessment of urine copper excretion in WD, its collection in children is difficult and it is not practical in large scale screening. Our previous attempts to develop a cut-off value of spot urine magnesium in the assessment of body magnesium status motivated us to develop a similar index for evaluation of copper excretion (11). These spot urine excretion indexes may have a role in triage of patients with suspected Wilson’s disease or even population screening of pre-symptomatic Wilson’s disease among preschool children.

To make such a spot urine index useful, we must first evaluate the reference intervals (RI) which are important in the interpretation of all laboratory test results. Although large scale programs like CALIPER have been set up to better define RI in the paediatric age group, they are confined to analytes in serum or blood samples (12). It is reasonable as they are more commonly encountered in routine laboratories.

In this article, we first examined the reference intervals spot urine copper excretion indexes among 153 preschool children. Then, data of urine excretion from prevalent paediatric WD patients were compared to these reference intervals to show the potential test utility of these biomarkers. The results are promising with good sensitivity, suggesting that WD screening by spot urine in preschool children is feasible.

## 2. SUBJECTS AND METHODS

The reference range of spot urine copper excretion in preschool children was established from 153 archival urine samples. These archival samples had been collected in a previous community-based studies to establish reference standards of forced expiratory indices in Hong Kong Chinese preschool children (13) and to investigate the association between urinary metal excretion and lung function in these subjects (14). In brief, spot urine samples had been collected from preschool children (age range of 3 to 7 years old) on-site at the participating kindergartens. Spot urine samples were properly collected into acid-washed bottles which were specially prepared for trace element analysis to avoid contamination by the environment. The samples were transported back to our laboratory within 4 hours and were stored at a -80°C freezer until analysis in one batch.

Spot urine copper concentration was measured by inductively coupled plasma-mass spectrometry (ICP-MS) 7700 with octopole reaction system, Agilent Technologies. The measurement procedure included mixing an internal standard containing rhodium with the urine sample. Urine samples were warmed to room temperature before dilution with pre-treatment standard containing solution. After vortex, the samples were loaded into the ICPMS 7700 Analyser together with standard calibrators and QC samples. ICPMS is a powerful analyser for measurement of multiple elements. In-house assay has a detection limit of 0.02 µmol/L for copper. The mean concentration of Low QC sample of 0.66 µmol/L had an analytical CV% of 5.4%. Urine creatinine was measured by a modified Jaffe reaction on autoanalyers of Roche Diagnostics. Urine osmolality was measured by freezing point depression on automated osmometer.

Six WD patients who were diagnosed before the age of 11 and had been followed up in outpatient clinic were invited to participate. Their diagnosis had been confirmed by mutation analysis of the ATP7B gene (10). 2-3 spot urine samples of the same day in addition to a next day standard 24-hour urine sample for copper excretion monitoring were collected. Patients’ data were subdivided according to the time of spot urine collection, 4 of them had samples collected within 1 year of diagnosis and treatment (labelled as WD patients). They and other patients also collected spot urine after being treated for longer than one year (labelled as WD patients on treatment). Those patients after treatment for more than one year might have a reduction of copper load and urine copper excretion. Some patients collected urine at multiple occasions more than one year apart and their samples were labelled accordingly. Informed consent will be obtained from their parents. The study has been approved by institutional clinical research ethic committee.

The regression between spot urine copper concentrations were examined against spot urine creatinine and osmolality. Regression lines and confidence intervals of regression lines were plotted for controls. In addition, the prediction region covering the predicted distribution of 95% control data points were shown (blue rectangles in scatter plots, e.g. figure 2). Difference in spot urine indexes are compared between WD and control by non-parametric group-wise statistics (Wilcoxon test). Statistical significance was defined by type I error of 0.05.

**Figure 1A:**
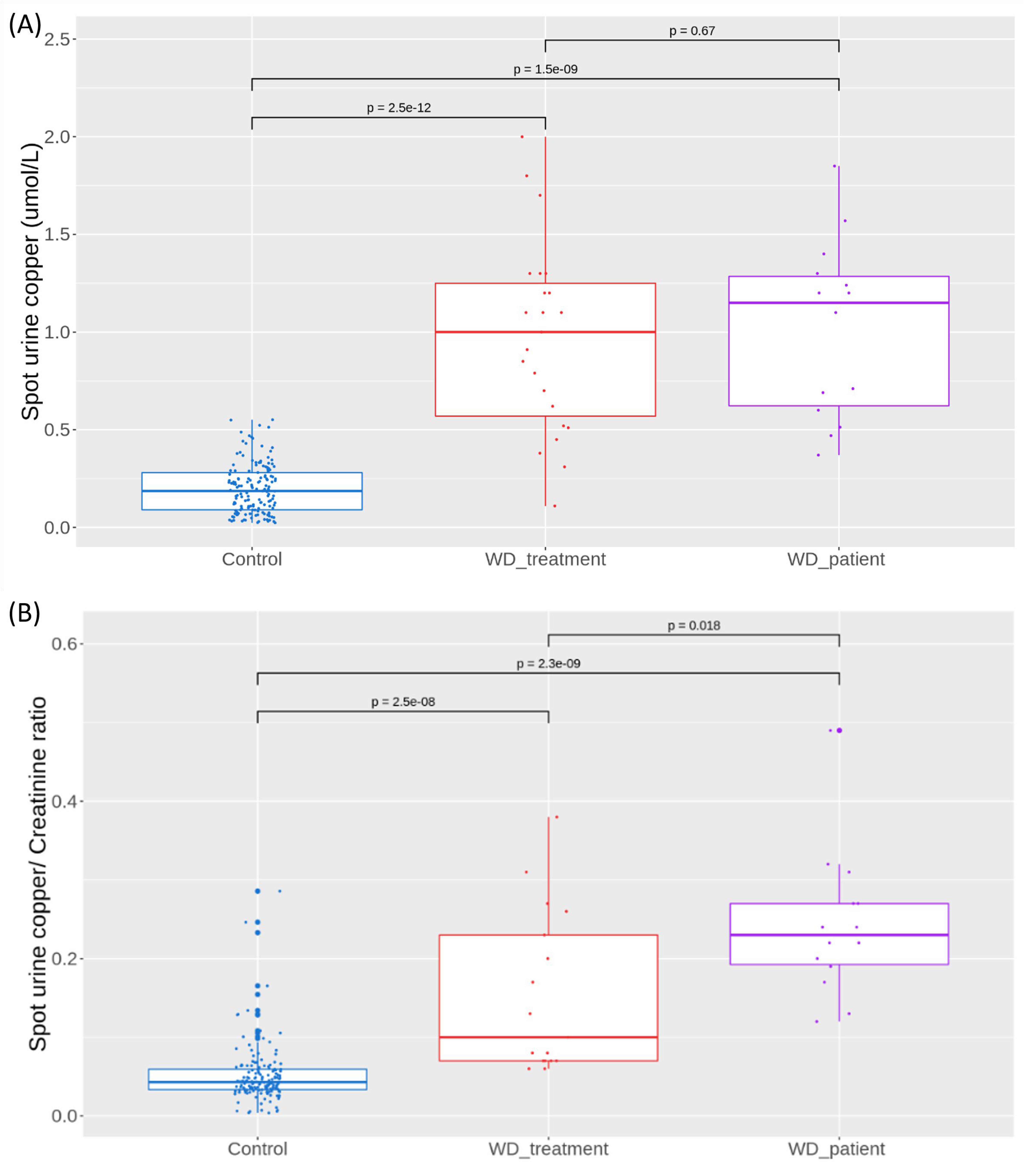
Figure 1A shows distribution of spot urine copper concentration of control (N=153), WD patients at diagnosis (WD_patient) and WD after more than 1 year of treatment (WD_treatment). The mean value of control was 0.2 µmol/L and few subjects had values above 0.5 µmol/L while most of WD patients had concentration higher than 0.5 µmol/L. Figure 1B: Figure 1B shows the distribution of urine copper to creatinine ratios (µmol/mmol) among three groups of subjects. All WD patients at diagnosis had ratio above 0.1. The value of control group was significantly lower than the two patients’ groups (p values <1e-7).

**Figure 2A:**
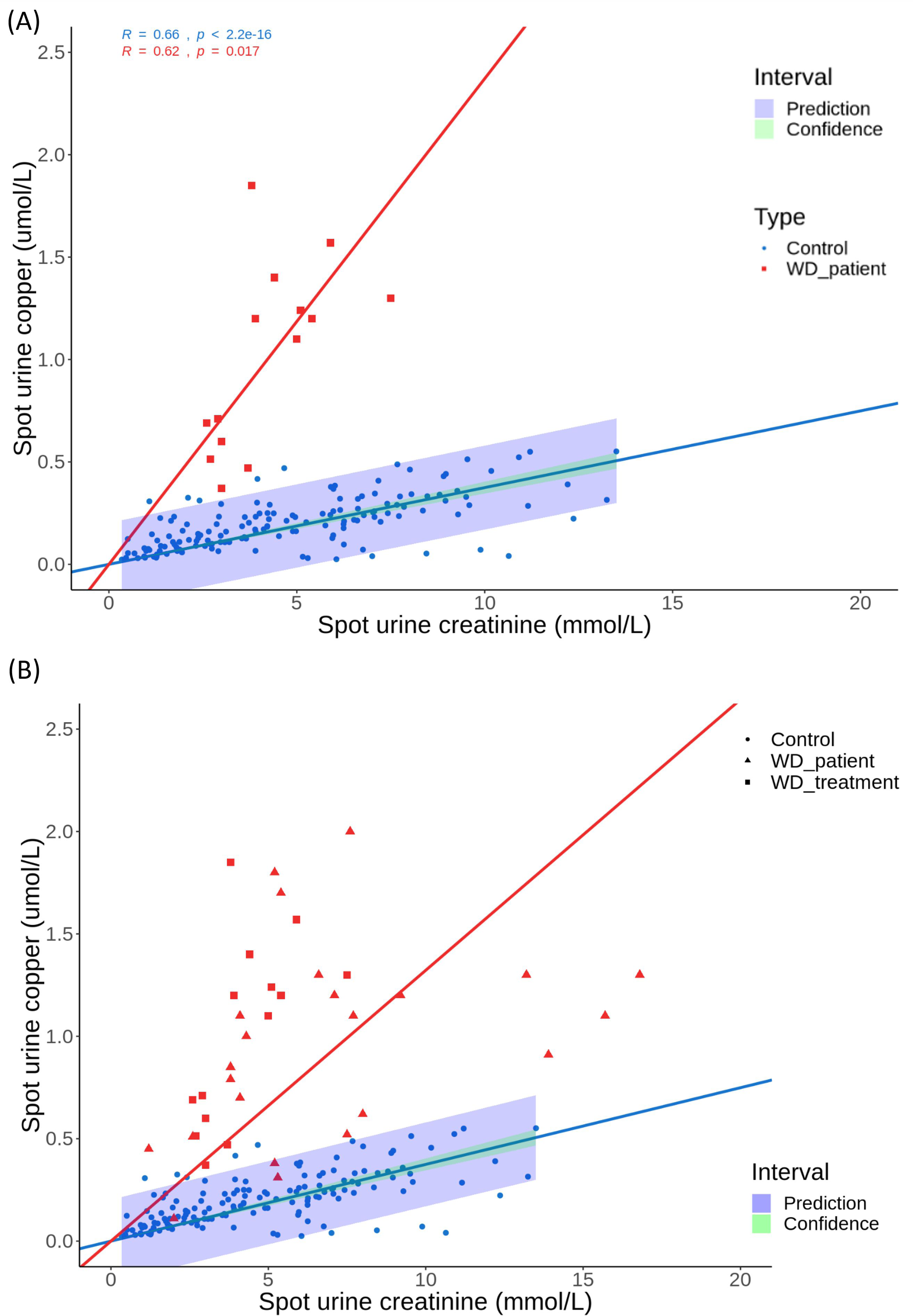
Figure 2A shows a strong correlation between urine copper and urine creatinine concentration in controls. The slope is much steeper in the WD patients at diagnosis. The blue rectangular band shows the predicted distribution area of control samples. Figure 2B: Figure 2B includes all WD patients. WD patients after treatment (WD_treatment, triangle symbol) had reduced copper to creatinine ratio and three samples were inside the predicted area of control distribution.

## 3. RESULTS

### Spot urine copper excretion indexes in preschool children (Table 1)

Among the 153 preschool children older than 3 years, the spot urine copper concentration had a median 0.19 µmol/L with inter-quartile values from 0.09 to 0.28 µmol/L. Only 4 children (3%) had a urine copper concentration exceeding 0.5 µmol/L. These values were 0.51, 0.52, 0.54 and 0.55 µmol/L. Although there was a mild age effect on spot urine copper concentration (increase by 0.026 µmol/L per year, p=0.02), the effect of age was corrected after normalization by creatinine or osmolality supporting the use of these ratios or bivariate interpretation (Supplemental Figure 1).

**Table 1:**
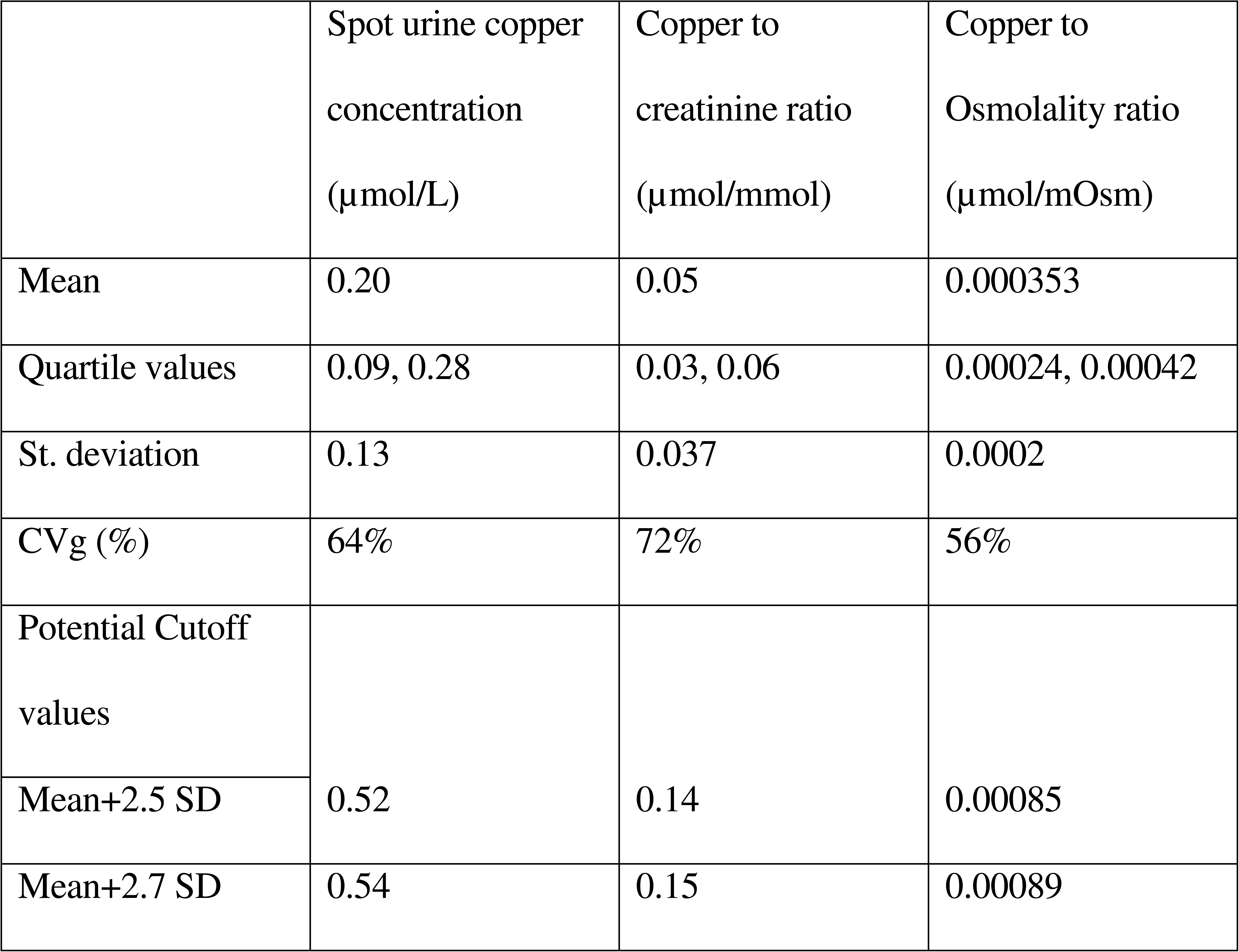
Distribution of spot urine copper excretion in preschool children between 3 and 10 years old (N=153). CVg stands for inter-individual variation in controls. Urine Copper to osmolality has the lowest CVg and the proposed cut-off values 0.00085 µmol/mOsm represented the mean + 2.5 SD value.

| | Spot urine copper concentration<br>($\mu\text{mol/L}$) | Copper to creatinine ratio<br>($\mu\text{mol/mmol}$) | Copper to Osmolality ratio<br>($\mu\text{mol/mOsm}$) |
| --- | --- | --- | --- |
| Mean | 0.20 | 0.05 | 0.000353 |
| Quartile values | 0.09, 0.28 | 0.03, 0.06 | 0.00024, 0.00042 |
| St. deviation | 0.13 | 0.037 | 0.0002 |
| CVg (%) | 64% | 72% | 56% |
| Potential Cutoff values |  |  |  |
| Mean+2.5 SD | 0.52 | 0.14 | 0.00085 |
| Mean+2.7 SD | 0.54 | 0.15 | 0.00089 |

The two normalised indexes were investigated, namely copper to creatinine ratio and copper to osmolality ratio and their mean values were 0.05 µmol/mmol and 0.000353 µmol/mOsm, respectively. Copper to osmolality ratio had the least biological variance (inter-individual variation) and CVg was 56% which was comparable to other spot urine tests, for example, urine microalbumin test for Diabetes nephropathy. On the other hand, copper to creatinine ratio had the highest CVg.

On the other hand, spot urine copper excretion was much higher in the patient groups irrespective of duration of treatment. Among 37 spot urine samples (with creatinine results) collected from all 6 WD patients, the median copper concentration was 1.10 µmol/L with inter-quartile values from 0.60 to 1.30 µmol/L. There were 6 samples (19%) with spot urine copper below 0.5 µmol/L (Figure 1A). Most of them (4 out of 6) were collected from patients whom received more than one year of treatment so these urine samples were categorised into the subgroups in subsequent analysis.

### Spot urine copper correlated with urine creatinine in control preschool children

It is a common practice to normalised spot urine excretion by creatinine. Indeed, spot urine copper concentration showed a strong correlation with spot urine creatinine concentration among controls (Pearson r=0.62, p≤2.2×10^-16^, Figure 2a). The predicted range of 95% distribution of individual data points (Prediction interval) are shown as blue rectangles in Figure 2a and 2b. The predicted range shows a good match with the actual distribution of data points. Only three control urine samples had urine copper concentration above the prediction range and another 5 samples were below, which represented ∼5% (Figure 2a). Increased spot urine copper concentration among WD patients were clearly separated from the predicted control range. The results indicate that bivariate analysis of spot urine copper excretion or copper/creatinine ratio should be informative for diagnosis of WD in preschool children.

For patients after long term treatment, spot urine copper excretion tends to reduce and approach towards that in the control (red triangle symbols in Figure 2b). Three WD patient (after treatment) samples fell into the prediction range of control (Figure 2b). The 2 regression lines (one for control and one for WD patients) represent urine copper to creatinine ratios and difference in two slopes could clearly differentiate between patients and control. The median urine copper to creatinine ratio were 0.23 (WD patients at diagnosis) and 0.17 (WD patients after treatment), which were almost 6 folds and 4 folds higher to that of the controls, 0.04 (Figure 1B). Therefore, 0.1 may be a reasonable cut-off value for urine copper to creatinine ratio to demarcate WD patients from controls which was marked in the subsequent ROC plot (Figure 4). In addition, the spot urine may also be useful in monitoring patients’ responses after treatment.

**Figure 3A:**
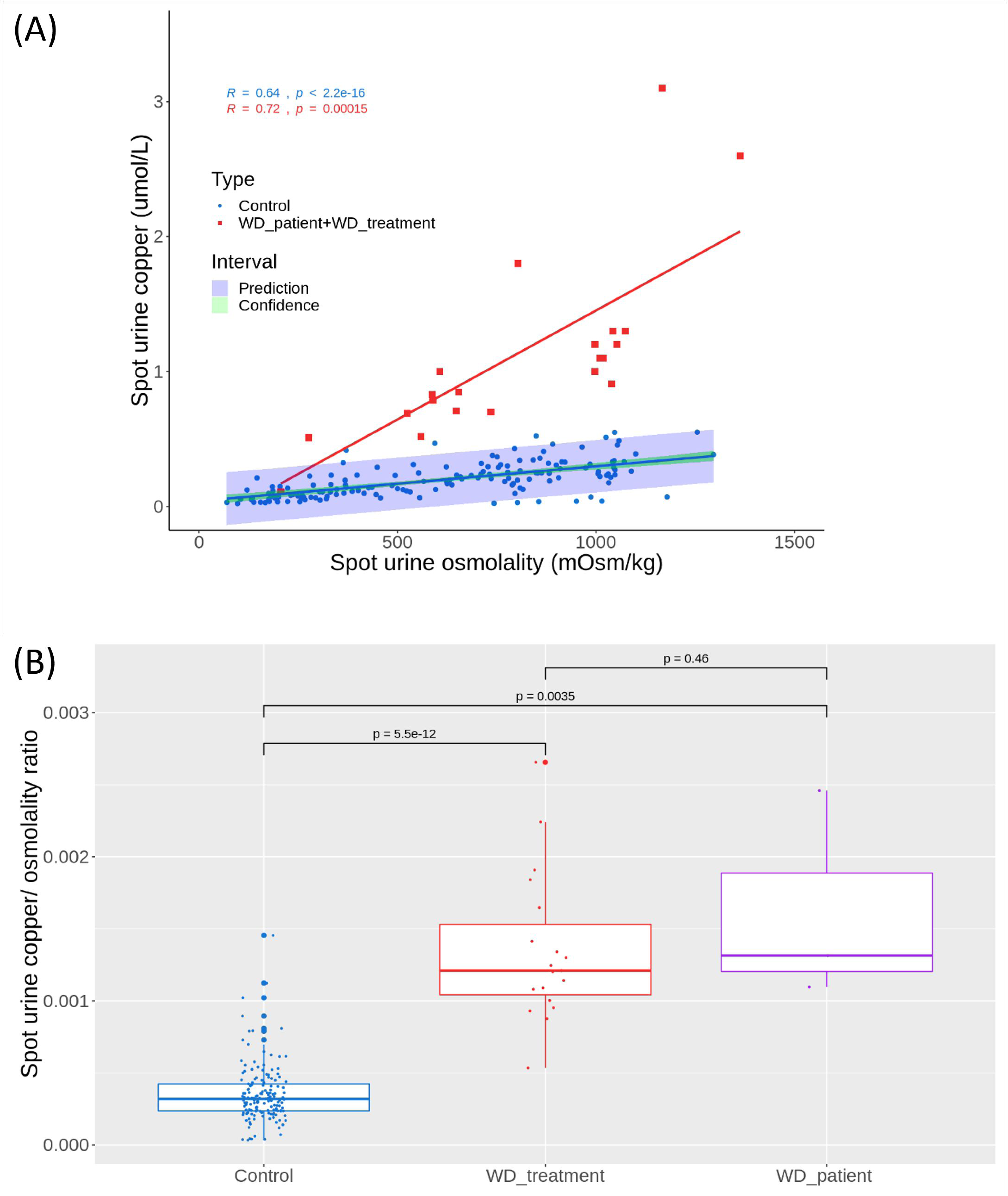
Figure 3A shows a strong correlation between urine copper and urine osmolality in control (R=0.64). The slope is much steeper in the WD patients. The blue rectangular band shows the predicted distribution area of control samples. Figure 3B: Figure 3B shows the distribution of urine copper to osmolality ratios (µmol/mOsm) among three groups of subjects. All WD patients at diagnosis had ratio above 0.00085. The value of control group was significantly lower than the two patients’ groups (p values <1e-12).

**Figure 4:**
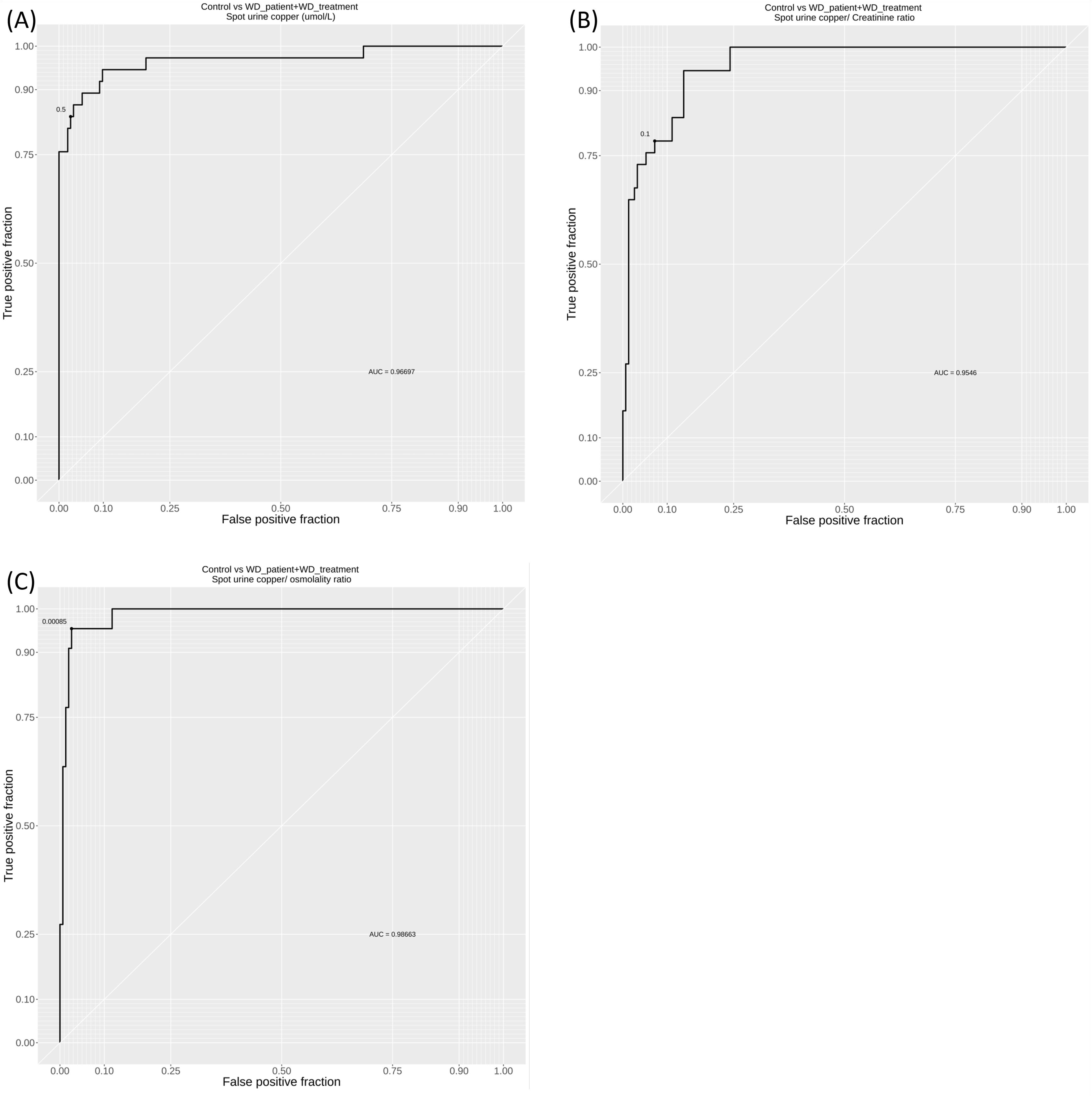
Figure 4 shows three ROC curves for diagnosis of WD using (A) spot urine copper, (B) copper to creatinine and (C) copper to osmolality ratios and cutoff values used are 0.5 µmol/L, 0.1 µmol/mmol and 0.00085 µmol/mOsm, respectively. While all parameters had area under curve greater than 0.95, copper to osmolality ratio had the best AUC of 0.986.

### Spot urine to osmolality ratio is the best biomarker

Likewise, spot urine copper concentration showed a highly significant correlation with spot urine osmolality among controls (Pearson r=0.6, p=<2.2×10^-16^, Figure 3A). In the scatter plot Figure 3A, there is a good demarcation between patients and controls. As only 3 data points were available from WD patient at diagnosis, all WD samples (N=22) were analysed together. There was only one WD sample that was located in the predicted range of control data. This samples were one of the more dilute urine samples collected from a patient at osmolality of 206 mOsmol/kg. When the distribution of the copper to osmolality ratio (Figure 3B) were analysed, the control group had a median of 0.00032 and a quartile range from 0.00024 to 0.00043. Meanwhile, the median in WD patients were 0.0012 with a quartile range from 0.001 to 0.0016. A cut-off value of 0.00085 could be used to differentiate WD from controls.

### ROC analysis of various spot urine parameters

Cut-off values for spot urine copper concentration, copper to creatinine ratio and copper to osmolality ratio at 0.5 µmol/L, 0.1 µmol/mmol and 0.00085 µmol/mOsmol (the cut-off values are 32 µg/L, 56 µg/g creatinine and 0.054 µg/mOsmol, respectively, in conventional units) have potential application in differentiation of WD patients.

In fact, both copper concentration and the copper to creatinine ratio can easily differentiate them. ROC analysis of all indexes had area-under-curve (AUC) values of greater than 0.95 and over 90% sensitivity in identification of WD from health preschool children control (Figure 4).

The best index was spot urine copper to osmolality ratio, which had the highest AUC of 0.98. Both sensitivity and specificity were better than 0.95 at cut-off value of 0.00085. Therefore, it is likely that such bivariate classifier will be used in future implementations.

For example, there was an overlap if the urine sample was dilute urine (at the low end of creatinine concentration). Therefore, low spot urine osmolality or creatinine concentration could be used as an additional filter to call for repeat. For example, spot urine osmolality lower than 500 or creatinine concentration lower than 1.5 mmol/L will be used as criteria to call for repeat samples.

## 4. DISCUSSION

WD is among the most prevalent IMD in Chinese and many ethnic groups. A population scale screening program is highly desirable for making early diagnosis. Furthermore, the recent use of oral zinc in the treatment of early diagnosed cases (before 10 years old) is highly promising with little side effect (15). Together with the natural history of WD, we propose to utilities this new screening window targeting preschool children or school children before 10 years-of-age. This new screening window has the advantage that there is a substantial and detectable body copper load at this age, while the patients are still largely asymptomatic and will response well to treatment. This study is carried out to investigate if screening of preschool children is feasible by firstly understanding the reference excretion of copper in this age group. Here we report that various spot urine copper excretion parameters are informative in differentiating WD from preschool controls, which will make wide scale screening feasible.

### In search of a suitable biomarker for large scale screening of WD

Timing of screening for WD is very important. Many previous attempts with limited success had targeted to screen WD at newborn (5). However, such approach is restricted by the limited exposure to copper of the foetus before birth resulting a small body copper load at birth. Here, we propose a novel screening strategy to make use of the fairly long presymptomatic phase of WD to screen at preschool age or before 10 years-of-age. Such flexibility also allows better integration into the local children health surveillance program. Starting with a non-invasive spot urine biomarker, large scale screening is logistically feasible and well received by both children and parents.

Our long-term interest in nutritional research and biomarker studies provided us an insight into the application of spot urine as biomarkers (11,16). In the past, we pioneered the use of spot urine magnesium as a biomarker for assessment of body status of magnesium (11) and it had been used in both clinical and research settings (17,18). There were other groups which carried out pilot studies in the use of spot urine in assessment of body status or dietary intake of sodium (19–21), magnesium and other trace elements (22,23). Spot urine collection is a common practice in the assessment of renal function in children. For example, spot urine protein to creatinine and urine albumin to creatinine ratios are commonly used in paediatric nephrology. They serve as reliable biomarkers in particularly this younger age group as the variation of body anthropometry is less in extent than in adults. Furthermore, the inter-individual biological variations in daily creatinine production and excretion are more stable (24). In a latest review, Armer et al also suggested a potential use of spot urine copper indexes in WD (25). However, the differences and test performance of spot urine copper indexes between WD and control have not been fully investigated. Furthermore, the evaluation of these variations in the paediatric age group is also lacking. Our study fills in this knowledge gap and supports the potential of use of spot urine copper index (in particular spot urine to osmolality ratio) in the screening of WD among preschool children.

With the ultimate goal of identifying a screening biomarker for WD of high sensitivity and specificity, both spot urine copper to creatinine and urine to osmolality ratios showed a huge potential in this study. While all spot urine indexes were informative, urine to osmolality may have an edge over other 2 parameters. An example strategy using bivariate screening approach is shown in Figure 5. Spot urine biomarkers have several obvious advantages over other blood markers. Firstly, it is non-invasive and large scale screening application can be easily implemented and compliance is not a problem. Secondly, repeat sample or a multiple sample collection protocol could be arranged. Advances in molecular genetics nowadays enable the definite diagnosis of WD to be made robustly (26). The presence of locally prevalent mutations also enhances the diagnostic efficiency. Presymptomatic WD patients picked up by spot urine screening can be followed by a molecular test to make the definite diagnosis.

**Figure 5:**
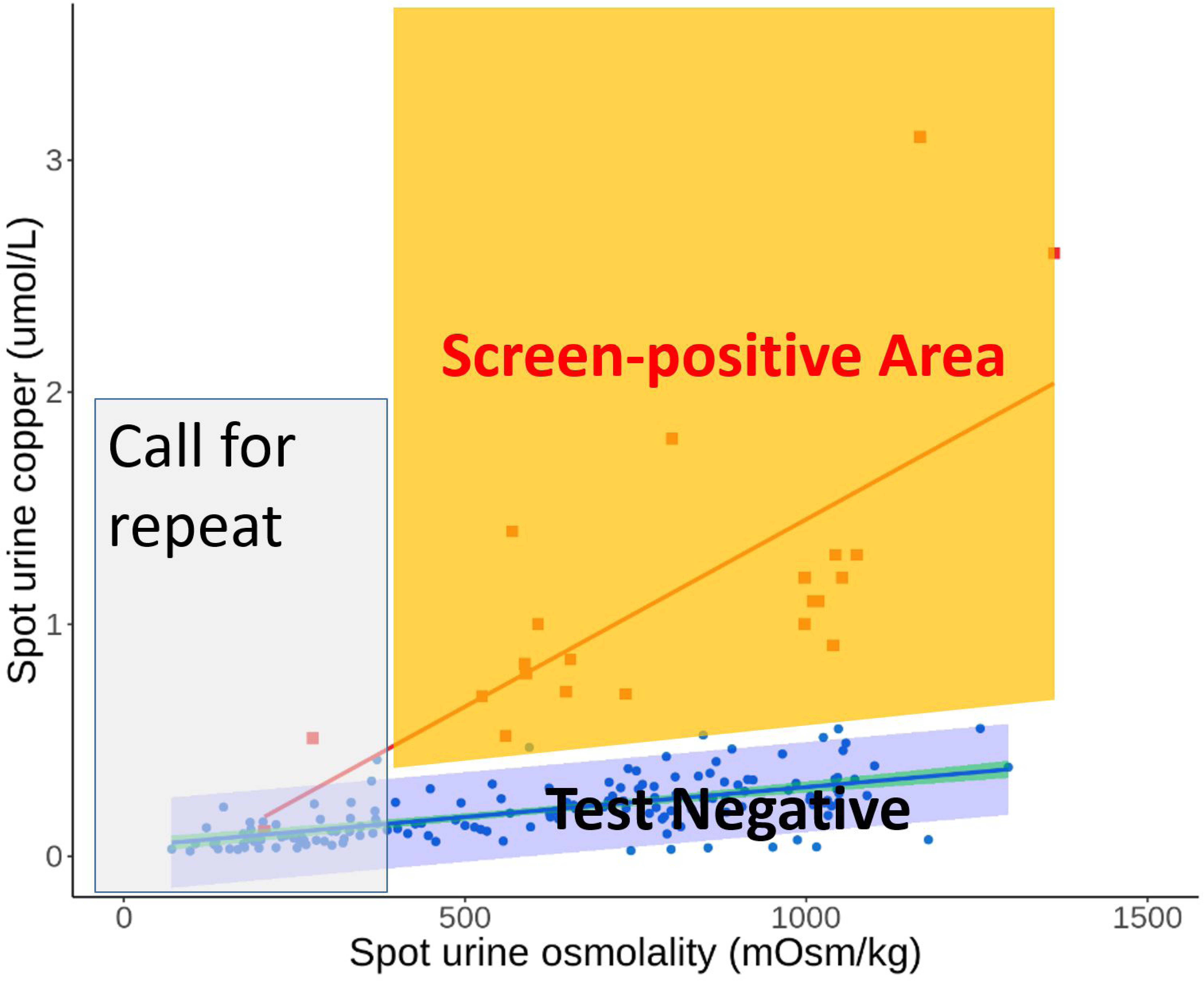
Figure 5 illustrates a potential screening strategy using bivariate data of spot urine copper and osmolality. Samples of low osmolality will be taken as non-diagnostic and subjects will be called to repeat. Subjects with samples inside the screen-positive area will be followed up with additional definitive tests.

While there is a long-held myth that urine copper excretion is highly variable (3), we showed that it was not the case in the control preschool children. In fact, urine excretion of magnesium and copper had the least biological variation among all trace metals, in terms of intra-individual variation (27). The data suggests that spot urine concentration of copper correlates with the average daily concentration in a 24-hour urine. This property of low biological variation provides the explanation that spot urine excretion is a reliable biomarker for selected elements like magnesium and copper. In the past, many confounding factors of urine copper excretion had been described, e.g. kidney disease, acute liver failure, hepatitis which were commonly listed as differential causes of elevated urine copper excretion. However, many of them are only found in the hospital setting but do not apply if urine is collected from presymptomatic (healthy) subjects. Therefore, they do not come into the picture in the setting of screening for presysmptomatic WD among healthy children.

There is a normal range of excretion in relation to creatinine or osmolality in the spot urine sample. These parameters differentiate early WD patients from controls fairly robustly with AUC greater than 0.95. On the other hand, such differentiation may be blurred in symptomatic WD patients who also had proteinuria or patients with other causes of proteinuria, as spot urine copper assessment is no longer valid for assessment of body copper status. Therefore, it is advisable to perform a proteinuria assessment together with spot urine copper as a screening package or subsequent follow-up test. Any samples with significant proteinuria will invalid the screening.

### Limitation of the study

This is the first attempt to use a bivariate analysis of spot urine for screening of WD in preschool children. We had data from only 6 WD patients in this age range. Although our results are encouraging, more data from childhood WD patients are required to fully understand the biological variation of spot urine copper excretion in early WD. Future multi-centre studies will be required to gather data from more patients.

## List of abbreviations

ROC: Receiver operator curve
AUC: area under curve
WD: Wilson’s disease
IMD: inherited metabolic diseases

