## Supplemental Figure 1 for "Reference intervals of spot urine copper excretion in preschool children and potential application in pre-symptomatic screening of Wilson’s disease"

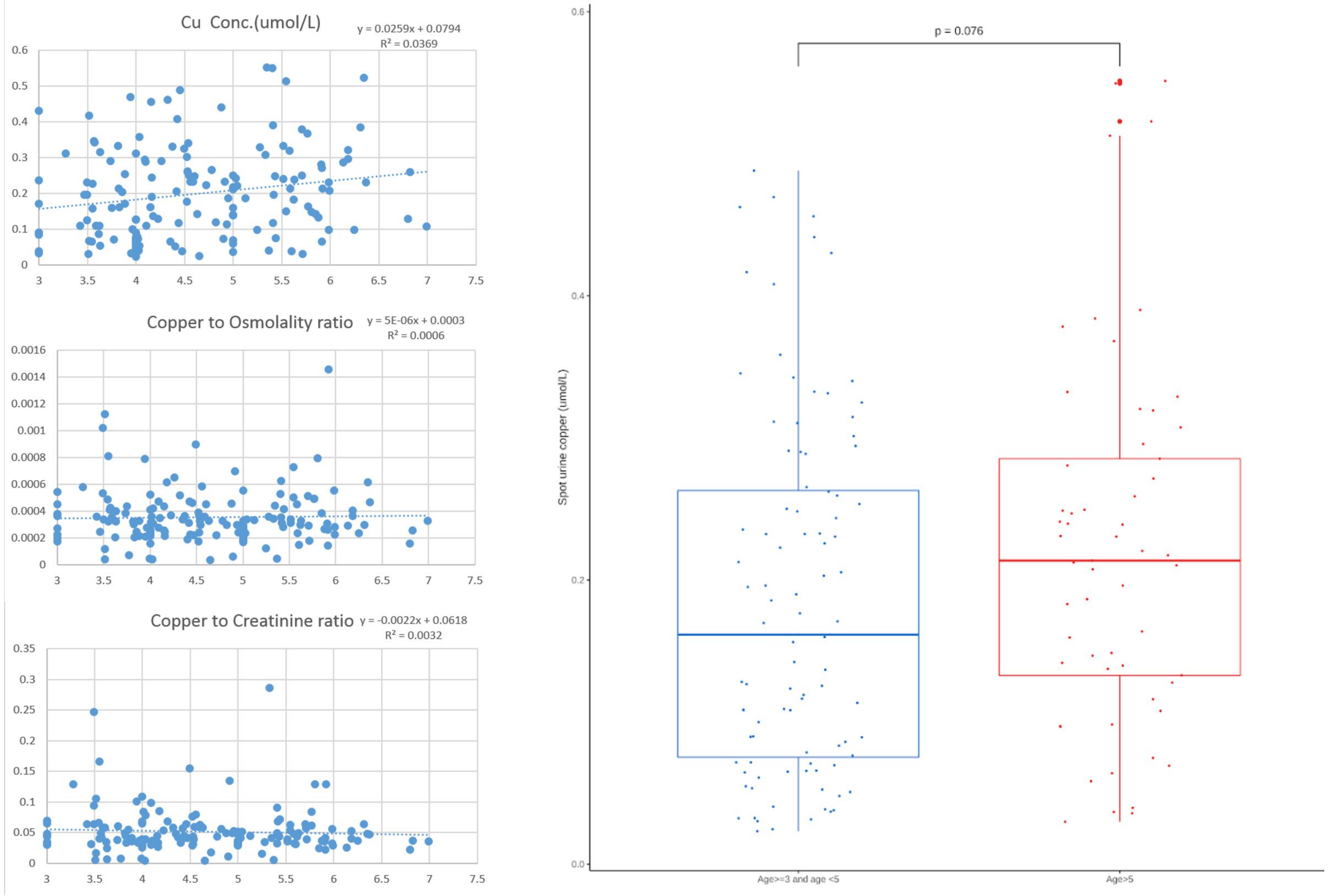


**Supplemental Figure 1:**

Figures show age effect on the spot urine copper excretion indexes. Y axis represent spot urine copper concentration, copper to creatinine ratio and copper to osmolality ratios. Although there was a mild age effect on spot urine copper concentration (slope=0.026, $p=0.02$), the effect was small as the difference was not significant after dividing the controls into 2 groups of below and above 5-year-old (Box plot, $p>0.05$). Furthermore, the effect of age was not observed after normalization by either Osmolality or Creatinine.
